# Humans account for cognitive costs when finding shortcuts: An information-theoretic analysis of navigation

**DOI:** 10.1101/2022.08.06.503020

**Authors:** Gian Luca Lancia, Mattia Eluchans, Marco D’Alessandro, Hugo J. Spiers, Giovanni Pezzulo

**Author notes:** Contact info for correspondence:* Dr. Giovanni Pezzulo, ISTC-CNR, Via S. Martino della Battaglia, 44, 00185 Roma RM, Italia.

## Abstract

When faced with navigating back somewhere we have been before we might either retrace our steps or seek a shorter path. Both choices have costs. Here, we ask whether it is possible to characterize formally the choice of navigational plans as a *bounded rational* process that trades off the quality of the plan (e.g., its length) and the cognitive cost required to find and implement it. We analyze the navigation strategies of two groups of people that are firstly trained to follow a “default policy” taking a route in a virtual maze and then asked to navigate to various known goal destinations, either in the way they want (“Go To Goal”) or by taking novel shortcuts (“Take Shortcut”). We address these wayfinding problems using InfoRL: an information-theoretic approach that formalizes the cognitive cost of devising a navigational plan, as the informational cost to deviate from a well-learned route (the “default policy”). In InfoRL, optimality refers to finding the best trade-off between route length and the amount of control information required to find it. We report five main findings. First, the navigational strategies automatically identified by InfoRL correspond closely to different routes (optimal or suboptimal) in the virtual reality map, which were annotated by hand in previous research. Second, people deliberate more in places where the value of investing cognitive resources (i.e., relevant goal information) is greater. Third, compared to the group of people who receive the “Go To Goal” instruction, those who receive the “Take Shortcut” instruction find shorter but less optimal solutions, reflecting the intrinsic difficulty of finding optimal shortcuts. Fourth, those who receive the “Go To Goal” instruction modulate flexibly their cognitive resources, depending on the benefits of finding the shortcut. Finally, we found a surprising amount of variability in the choice of navigational strategies and resource investment across participants. Taken together, these results illustrate the benefits of using InfoRL to address navigational planning problems from a bounded rational perspective.

## Introduction

Navigating a city or other everyday environments can require a myriad of decisions. These can be as simple as choosing to follow the usual route we have taken before, or as complex as devising a novel route through a vast array of possible paths to reach a novel goal, i.e., wayfinding. Recent experiments are increasingly shedding light on the variety of navigational strategies that we use to find our way in real-world settings (1–3) and virtual environments (4– 9), as well as their neural underpinnings (10,11). However, an integrative computational perspective is still missing.

A standard assumption is that navigational planning requires a form of cognitive *tree search* over a mental map (12–14). Tree search implies that people try out all (or many of) the possible alternative routes and finally select the shortest one. However, extensive tree search is not feasible, except in very simple situations. Therefore, it has been proposed that people might adopt various heuristics to alleviate the burden of exhaustive search (15). One such heuristic is pruning, or the idea that during the mental search, if a tree node is encountered that has a particularly low value, the whole branch of the tree is discarded (16). Another heuristic consists in sampling only a few routes, or multiple routes but only up to a certain depth, similar to Monte Carlo sampling methods in statistics (17,18). Finally, it has been proposed that people might use hierarchical forms of planning (or subgoaling) to split the problem into more manageable subproblems (1,10,11,19–21).

An alternative, less considered possibility is that navigational planning does not use tree search. Rather than searching the tree of possible routes, people might start from a “default plan” to follow a well-known (or habitual) route and then modify the plan when necessary. For example, if a person has to reach a shop close to her office, she can start by considering the usual route to the office and then adapt it (e.g., take a turn before reaching the office) to actually reach the shop, rather than the office. In this perspective, the “default plan” would act as an anchor for the planning problem and any deviation from it would require a (cognitive) cost. This idea has been formalized in terms of Information Reinforcement Learning (InfoRL) (22) and Linear Reinforcement Learning (23,24), but these theoretical formulations have not yet been applied to human navigational planning studies.

Here, we adopt the InfoRL formulation (22), which considers planning as a *bounded* rational process that entails a cost-benefit trade-off. The trade-off arises because the planner can select how much cognitive effort to invest to deviate from the “default plan” (where following the default plan is assumed to have zero cost) to improve the solution and potentially achieve a greater reward. In this navigation setting, we consider the reward to be proportional to (minus) the length (i.e., number of steps) of the route; hence, the shortest route is also the route having the highest reward. This formulation highlights that the benefit of investing a greater cognitive cost is that, by deviating from the default plan, one can potentially find a shorter route to the goal.

The decision to be made is therefore how much cognitive effort to invest. In InfoRL, the cognitive effort of planning can be called a complexity cost since in information terms, it can be measured as the amount of “control information” (in bits) to be processed to change the default plan (a *prior* in Bayesian parlance) to the final selected plan (a *posterior* in Bayesian parlance); see (22,25–28). In general, the lower the cost, the closer the selected plan to the “default plan”; whereas the greater the cost, the farther the selected plan from the default plan - and the greater the possibility to find a higher-reward (i.e., shorter) route.

The bounded rational InfoRL framework emphasizes that the goal of the planner is not, or not always, finding the shortest route (i.e., maximizing reward), but rather optimizing the cost-benefit trade-off between finding the shortest route (i.e., maximizing reward) and avoiding cognitive effort (i.e., minimizing control information). This trade-off could be different for different people. People might have different cognitive resource limits, which correspond to different thresholds on information costs in InfoRL. Furthermore, some people may prefer paying a greater cost to find a shorter path, whereas others might prefer paying a smaller cost and deviating less from the default plan. Therefore, there may be no single “optimal” solution to the cost-benefit trade-off, it depends on the ability of the person. However, while the solution for a given individual may vary, InfoRL permits calculating the maximum level of reward that is achievable for each level of effort investment. Specifically, this method considers what is the maximum reward that is achievable by an optimal agent that uses any given level of control information. The solution is not just one single optimal solution, but a *frontier* of optimal solutions (i.e., a trade-off curve, see Figure 1).

**Figure 1.**
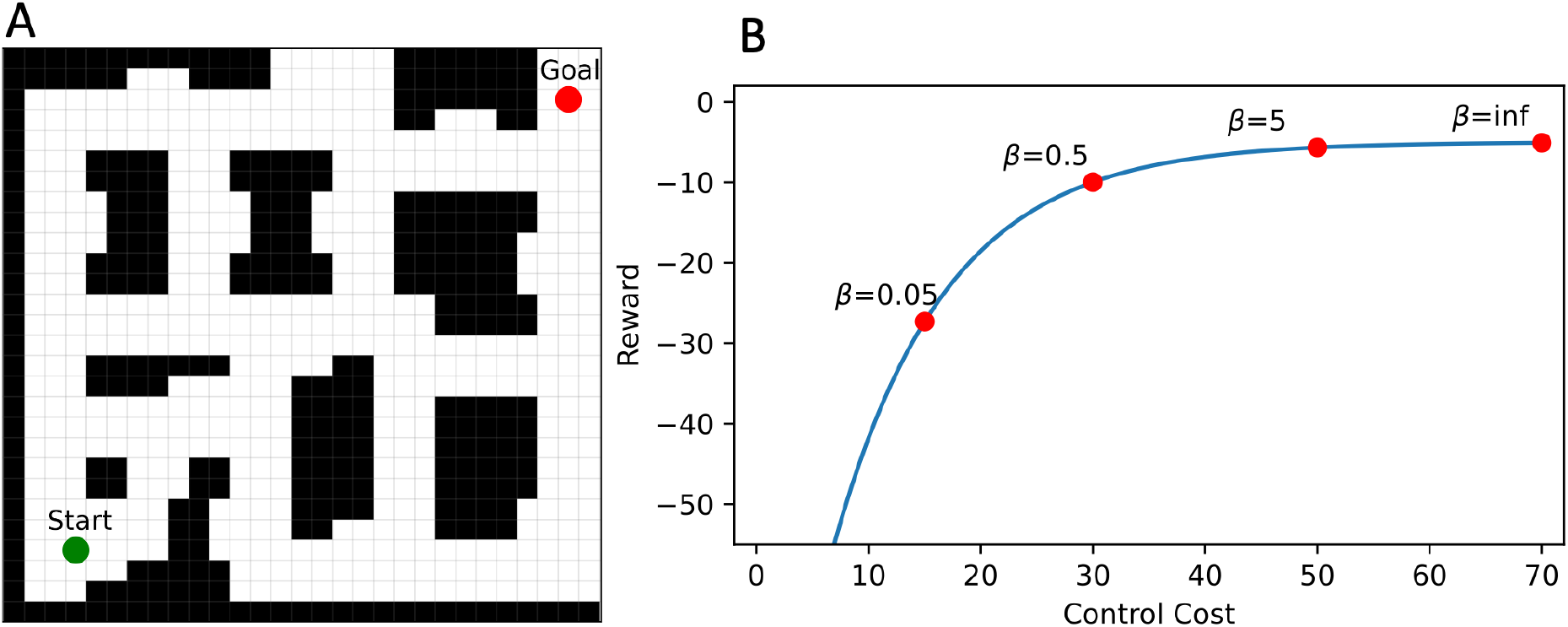
Theoretical trade-off curve for a spatial navigation problem. (A) An example navigation problem, where an agent starts at the green dot and needs to navigate to the red goal location, and the challenge is persisting along the path. (B) The blue curve represents a “frontier” of optimal solutions achieved by optimal RL agents (red dots). Each of the optimal RL agents is characterized by a different beta parameter, which regulates the amount of control cost that the RL agent invests (x axis) to obtain different levels of reward (y axis). Note that reward here is proportional to (minus) the length (i.e., number of steps) of their route; hence, the shortest route is also the route having the highest reward. The bottom-left part of the blue curve shows the maximum level of reward achievable by using the default policy (and hence zero control information). Please note that in this example (but not in our main analysis, see below), the default policy is a random policy - i.e., there is no cost associated with acting randomly. The top-right part of the blue curve shows the maximum level of reward achievable with the shortest route, which deviates the most from the default policy and hence requires the most control information. Note that the curve shown in the figure is only illustrative and inspired by (29); see below Figure 4 for the InfoRL curve characterizing our navigation problem.

The example shown in Figure 1 illustrates a simple case of spatial navigation and its analysis using InfoRL. Please note that this is a very simple example of wayfinding, which does not involve any choice between different paths, but rather just determining a path between a starting location and a goal location. Yet, in InfoRL, planning a path (or policy in RL terms) requires some information cost. InfoRL permits analyzing navigational planning strategies that vary along the two axes of reward (e.g., length of the solution path) and control information (e.g., bits) and that achieve different levels of optimality. In the Figure, the blue line represents the “frontier” of optimal solutions: a theoretical trade-off curve calculated by considering “simulated participants” (red dots) that are actually optimal RL agents that use different amounts of control information (controlled by a beta parameter). Each “simulated participant” is characterized by three parameters. The first parameter (which is directly observed, not inferred by InfoRL) is the amount of reward obtained in the task, such as the length of its path during a navigational problem (see the y axis). The second parameter is the amount of control information used to solve the task (see the x axis). This second parameter can be inferred using InfoRL, by considering how much the participant’s route differs from the default plan. The third parameter is the optimality of the participant’s solution, given the level of control information used. This third parameter corresponds to the distance of the participant’s location in the plot from the optimal frontier (i.e., the distance from the red dot and the blue line). The “simulated participants” shown in Figure 1 are all optimal agents and therefore they lie exactly on the blue curve. However, real participants could lie anywhere below (not above) the blue line. A participant who reaches the frontier and achieves the maximum possible reward, at any level of control information, makes an optimal use of its resources. Rather, a participant who achieves a lower level of reward than allowed from its investment is planning suboptimally. Therefore, not only can InfoRL help quantify the amount of control information that people invest when planning but also whether they use this investment optimally to obtain the maximum information gain.

In this article, we ask whether the InfoRL framework (22) can be used to characterize human navigational planning in a virtual reality setting. One precondition to use InfoRL to study human navigational planning is being able to specify a default policy, which corresponds to the policy that people would consider by default, without investing any cognitive effort. In theoretical studies, the default policy is often assumed to be a random policy; but this is unreasonable in real-life navigational planning, where people (1) and mice (30) show systematic, non-random regularities. In cases where the environment is familiar it is more likely that people would, by default, follow a well-practiced, habitual route, which can be reactivated without using extensive cognitive resources (27,31,32). Following this line of reasoning, we applied InfoRL to a virtual navigation task, in which the concept of a default policy could be mapped to a route that was extensively practiced prior to a wayfinding test (33,34).

In the task conducted by Boone et al. (33), participants were firstly trained to follow a “default route” several times through a maze-like environment (Figure 2). During this training phase, they encounter multiple landmarks along the way. Afterwards, participants were allocated to one of two groups and placed in a series of landmark locations and asked to reach known landmark locations. For one group they asked to go to a location in whichever way they preferred (first group, “go to goal” instruction). The second group were instructed to travel to the goal by taking a shortcut (second group, “take shortcut” instruction). Crucially, the participants could use a set of diverse strategies, as a function of their individual preferences and/or task instructions: they could reach the landmark by following the default route (which passes through all the landmarks, but does not follow the shortest path) or invest various levels of cognitive effort to plan alternative and potentially shorter routes. This task therefore allows us to test the extent to which InfoRL can identify and formalize the different strategies that people use during navigational planning and their associated levels of cognitive effort and optimality. Furthermore, this experiment also allows testing differences in strategy selection and optimality between groups of participants that received different (“go to goal” or “take shortcut”) instructions. Finally, with the maze layout we can test whether participants deliberate more in places where it is necessary to invest more information to make informed choices (19,35).

**Figure 2:**
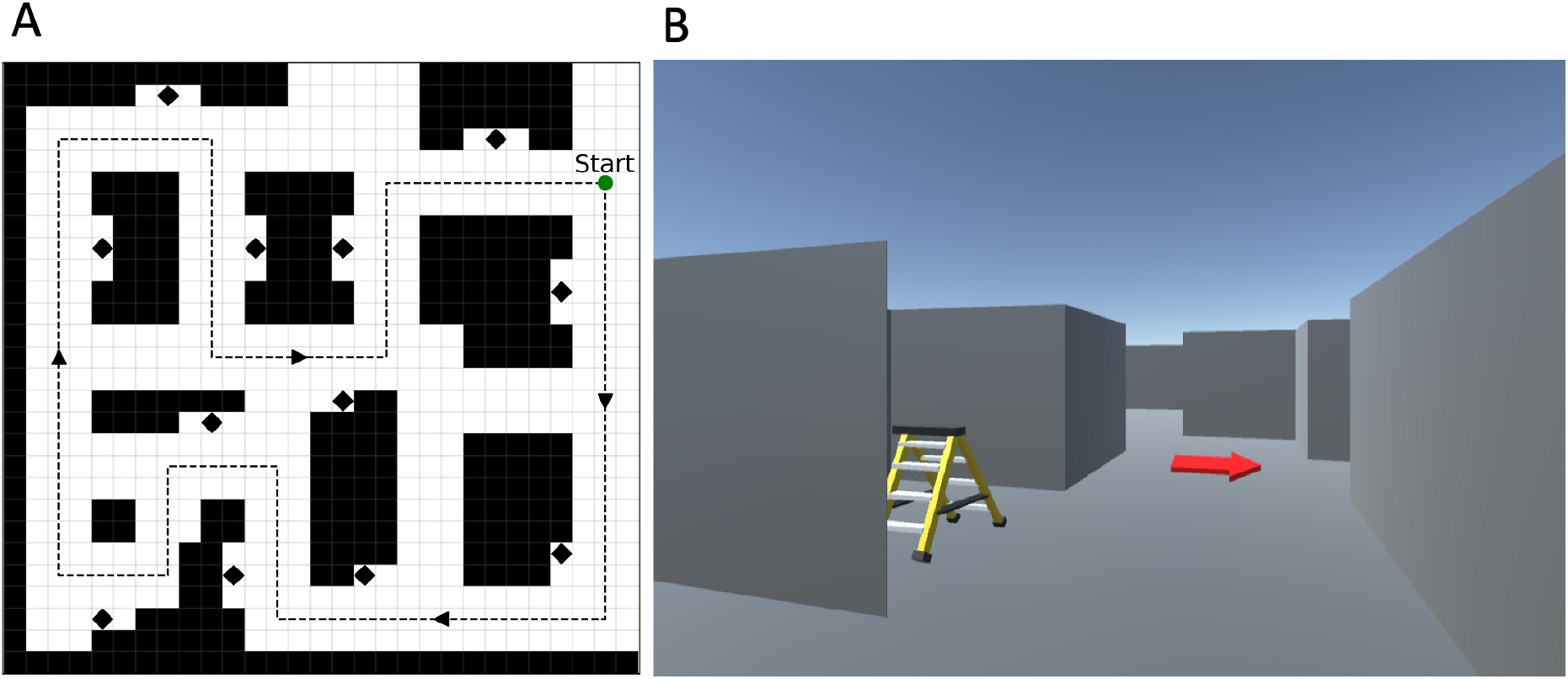
Experimental setup. (A) Top view of the map used in the experiment (not visible to the participants). The gray arrows show the stereotyped path used during the training phase and the black diamonds represent landmarks. (B) An example screenshot showing the participant’s view of the maze during the training phase. The red arrow indicates the training path. Figures redrawn from (33).

To preview our results, we report five main findings that illustrate the benefits of using InfoRL to study navigational planning and wayfinding. First, InfoRL permits characterizing formally different navigational strategies that correspond conveniently to different routes in the virtual maze. The strategies are also linked to path optimality (the ability to get the maximum reward from the amount of control information used) and suboptimality. Second, people deliberate more in places where the value of investing cognitive resources (i.e., relevant goal information) is greater. Third, while the group who receives the instruction to “Take Shortcut” seems naively more effective (purely considering path length), our analysis shows that not only do they invest more cognitive resources, but they also select optimal routes less often, compared to the group who receives the instruction to “Go to Goal”. This reflects the intrinsic difficulty of finding optimal shortcuts. Splitting participants by gender reveals that overall male participants invested more control information, but that both invested similar levels of increased control information in the “Take Shortcut” than the “Go To Goal” condition. Fourth, those who receive the “Go To Goal” instruction modulate flexibly their cognitive resource investment, depending on the benefits of finding the shortcut. Finally, we found that for most participants the amount of Control Information and Distance from the optimal curve was highly variable across trials. This implies that people do not stick to one strategy, but rather vary their strategy across trials.

## Results

### Experimental setup

The experimental setup is a first-person navigation game in a 3D virtual environment displayed on a flat screen, for which the data are freely available online (https://osf.io/ykxts/), see (33,36) and (34). The maze is relatively small, with 12 landmarks and 6 intersections (see Figure 2A for a map of the environment that was never shown to participants). The maze had gray walls and floor with landmarks standing out (see Figure 2B for an example first-person screenshot of what participants viewed).

The experiment of Boone et al. (33) is divided into two phases. In the first (training) phase, participants are forced to follow the stereotyped path indicated to them by red arrows in the environment (Figure 2B), for five times. The route taken is shown in a gray dotted line in Figure 2A. The training route passes 12 distinct nameable landmarks (a bicycle, a red stepladder, a TV, etc., see diamonds in Figure 2A), The portion of the environment marked with arrows is visible but not accessible during training.

In the second (experimental) phase, participants start from one of the landmark locations and are instructed to navigate to a goal landmark. This second phase is executed in 2 conditions, in 2 groups of participants: one group is instructed to reach the goal landmark (“Go To Goal” group) whereas the other group is instructed to reach the goal landmark location by using a shortcut (“Take Shortcut” group). The two groups of participants navigate in the map shown in Figure 2A. Participants execute 40 trials, each trial with a different starting position - goal landmark pair (please note that the original task was divided into 2 sessions, but here we combine them together for simplicity, as no differences in navigation efficiency were reported between the sessions). Each trial is recorded as a sequence of x and y coordinates, an angular coordinate representing the participant’s orientation in the map, and the respective timestamp. Trials which lasted over 40 seconds were not included in the dataset of (33).

For the data analyses, we discretized the map shown in Figure 2 into a graph composed of 66 connected nodes and which allows only 4 navigational actions (north, south, east, west); see Figure 3A. Figure 3B shows the default policy *ρ*(*a*|*s*), which includes the (deterministic) navigational actions performed by the participants during the training phase in the subset of nodes that were actually traversed (see the arrows in Figure 3B), and assigns a uniform probability to all the actions in the subset of nodes that were not actually traversed (see the yellow dots in Figure 3B). Please note that as in the experiment setup, a path under the default policy will visit all the goal landmarks in a finite number of steps.

**Figure 3:**
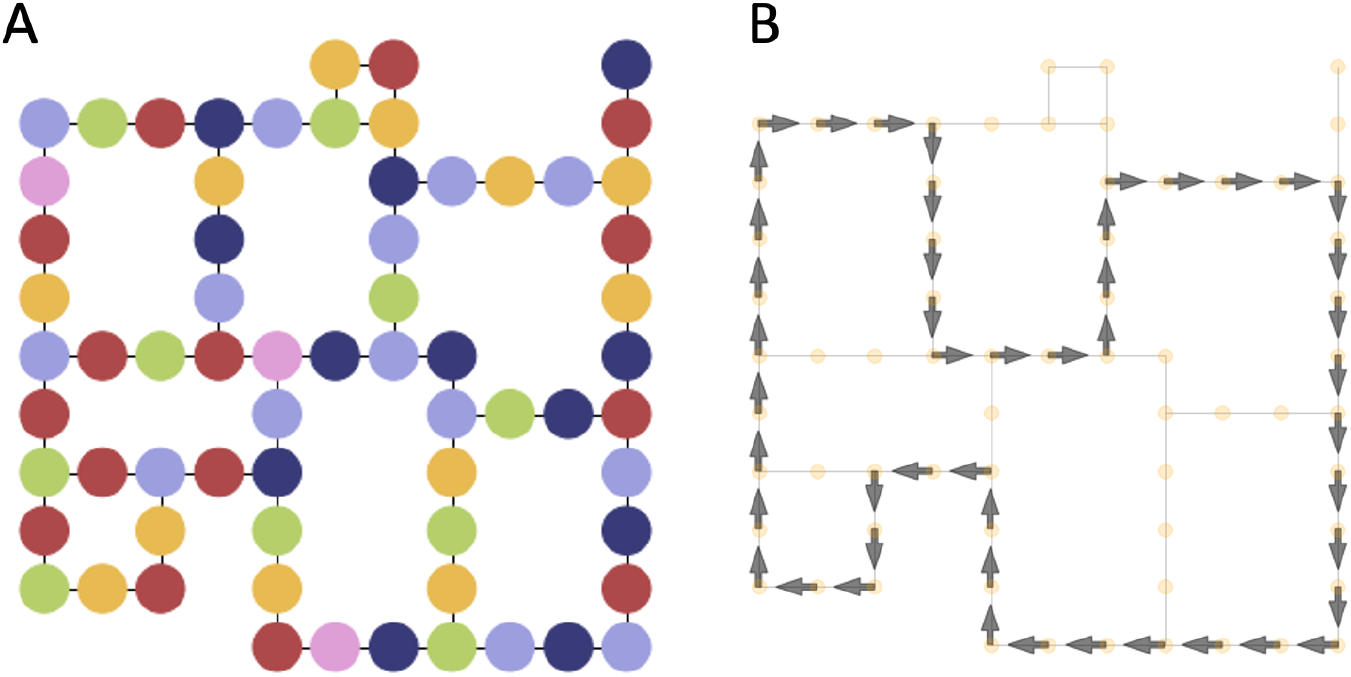
Data preprocessing. (A) We transformed the original map shown in Figure 2 into a graph of 66 connected nodes (which we coloured in arbitrary ways for ease of identification). Note that for each node, up to four (north, south, east, and west) actions are available. (B) Maze graph with arrows representing the default policy (the policy is uniformly distributed in the node without an arrow). See the main text for explanation.

### The Reward/Control Information trade-off characterizes navigational strategies and path (sub)optimality

We used the methods of (22) to characterize the participants’ paths formally, by calculating the “control cost” that each participant invests (where a low / high control cost means that the participant’s path is closer / farther from the “default policy”) and their “reward” (higher / lower reward indicates shorter / longer paths, with the shortest path being the highest reward path).

The curve shown in Figure 4A, left illustrates the optimal Reward / Control Information trade-off for one example task, whose start and goal locations are marked with red and green dots, respectively, in the small maps of Figure 4B-G. The solid black curve of Figure 4A shows the solutions of the optimal InfoRL agents that use different amounts of control information - and hence represents the theoretical limit on the amount of reward that participants can achieve, for each amount of control information. Each colored dot in Figure 4A corresponds to one participant and shows his or her placement in the Reward / Control Information plot.

**Figure 4:**
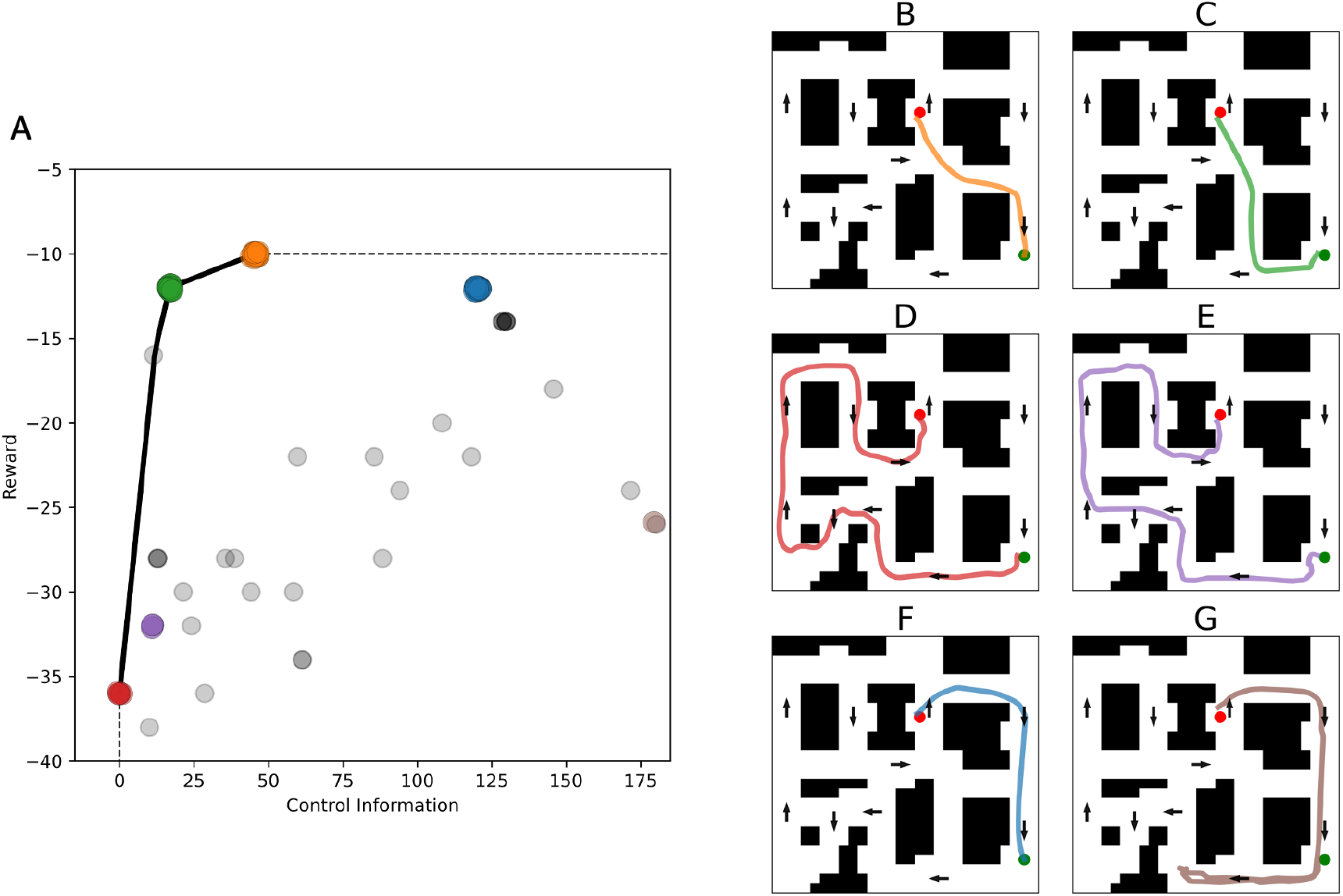
InfoRL analysis of navigational planning in virtual reality, applied to data from (33). (A) Reward-Control Information trade-off, for a single task. The solid black line shows the optimal trade-off function; the dashed gray line extends the optimal curve to separate the achievable area (lower right) from the unachievable area (top-left). Each colored dot corresponds to one participant in the task (we added random jitters to better show point clusters). Dot colors are matched to the paths shown in B-G. (B-G) Each figure shows a single participant’s path in the map, randomly extracted to represent the cluster. See the main text for explanation.

Two main findings are worth considering. First, several dots lie on the theoretical curve (orange, green, and red) or very close to it (purple and blue). These correspond to participants that behave optimally in the sense that they get the maximum reward from the amount of control information that they use. Rather, other dots are far from the theoretical limit and correspond to suboptimal participants that do not get the maximum reward from the resources they use.

Second, the dots having different colors in Figure 4A correspond to participants who select qualitatively different paths, see Figure 4B-G. This illustrates the fact that InfoRL permits identifying “clusters” of paths in the maze, optimal or suboptimal, with paths that are closer in the Reward-Control Information plot corresponding to similar paths on the maze. The paths shown in B-D correspond to three families of solutions (orange, green and red) that lie on the optimal curve. Specifically, the path shown in B (orange) is the optimal shortest trajectory; the path shown in C (green) is the optimal trajectory which requires the least Control Information; and the path shown in D (red) corresponds to a participant who follows the default policy and hence pays minimal control cost. The path shown in E (purple) is an off-curve, shorter but slightly more complex variant of the D path. The path shown in F (blue) follows the default policy, but in the reverse direction (this is why it has a high control cost). The path shown in G (brown) starts moving in one direction and then backtracks, consequently having both low Reward and high Control Information. This latter (G) path can be considered a longer (and hence suboptimal) version of the path shown in F. This can also be appreciated by noticing that in Figure 4A, the brown dots (one of which corresponds to the suboptimal solution G) lie diagonally but are more distant from the Reward / Control Information curve compared to the blue dots (one of which corresponds to path F). Similarly, many gray dots that lie below the Reward / Control Information curve can be considered as suboptimal (e.g., slightly longer) versions of the “optimal” paths that lie on the curve; please see the Supplementary Video V1 to better appreciate the different families of solutions.

In sum, InfoRL permits characterizing formally both the optimality of the participants’ solutions (as compared to optimal InfoRL agents) and the different families of paths / solutions that they selected for the navigational problem. Please note that Supp. Figure S1 shows the same plots for 20 pairs of start-goal locations, indicating the generality of the finding.

### Participants deliberate more at decision points where information demands for action selection are greater

InfoRL permits calculating the amount of Control Information that people invest to find a navigational plan and the benefits of such effort investment. However, InfoRL considers the information cost of selecting an entire path, not the information cost of single decision points during navigation.

During navigation, we cross various decision points that require investing more or less cognitive resources to select appropriate actions (or replan). The notion of Relevant Goal Information (RGI) (37) formalizes the information cost required to keep in mind (or ignore) the goal at any point during navigation. The RGI is important to the extent that selecting action in some places (e.g., at difficult decision points) requires processing information about the final goal, whereas selecting action in other places (e.g., along borders) does not -because in the latter places, the action to be selected would be the same, irrespective of the goal. The notion of RGI can be therefore considered an instantaneous measure of information cost during navigation - or an analogous of the notion of Control Information in InfoRL, but for single decision points.

We used the methods of (37,38) to calculate the RGI at each location in the map and asked whether participants spend more time deliberating and invest more cognitive resources in places of the maze where RGI is greater (see Figure 5A). As an index of deliberation, we used Vicarious Trial and Error (VTE) (39–41): a behavioral measure of how frequently a participant stops to look around (i.e., displays both low speed and high angular velocity) at each place of the maze. See the Methods section for details.

**Figure 5:**
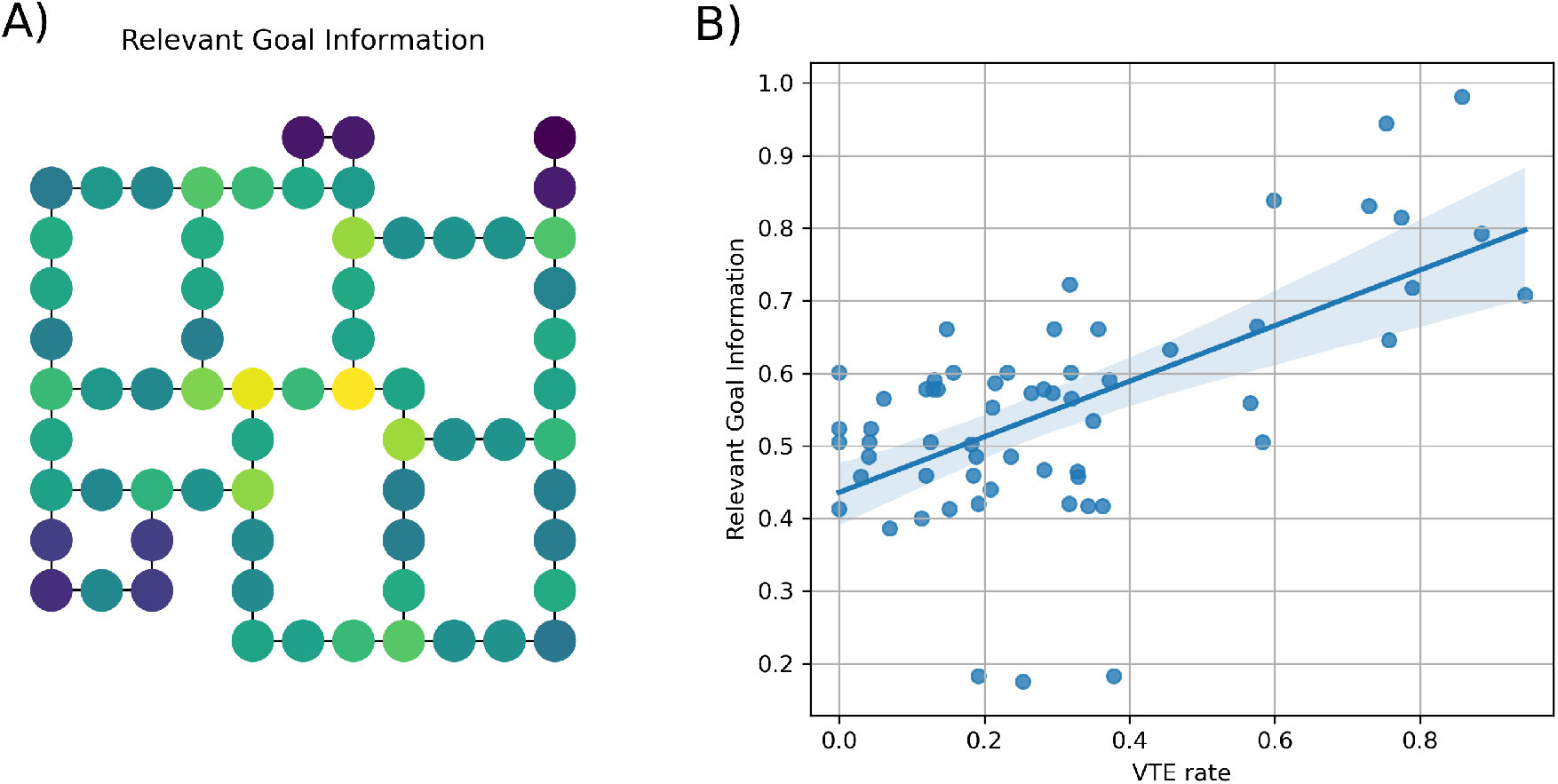
Deliberation as a function of information required for action selection. (A) Amount of Relevant Goal Information (blue represents low values, yellow high values) in different places of the map. (B) Correlation between Relevant Goal Information and rate Vicarious Trial and Error behavior; results are averaged across all trials for each state. Please note that we removed from the analysis four dead end states where the frequency of low-speed events was excessively high, regardless of decision processes. See the main text for explanation.

Figure 5B shows that RGI and VTE rates are highly correlated (Pearson ρ = 0.61, p < 0.01). This suggests that participants spend more time deliberating in places where action selection poses greater information demands. This result also indicates that planning and deliberation are continuous processes that occur during a journey (42,43), rather than processes that are completed before starting the navigation. Converging evidence for this latter point comes from the fact that we did not find any correlation between initiation time (i.e., the time before participants start moving) and planning difficulty or information cost. Rather than planning extensively before starting the navigation, participants started moving with little delay (mean 1.95 *s, SE* = 0.02) and plausibly planned (and replanned) all along the way, consistent with real-world reports of navigating (44).

Our control analyses (reported in the Supplementary Materials) show that VTE rates are also correlated with a graph-theoretical index widely explored in navigation studies: betweenness centrality (Pearson ρ = 0.64), which is a measure of how connected is a node to the rest of the graph, and is defined for a node in the graph as the fraction of shortest paths connecting two other nodes that pass through it. The correlations between RGI and VTE and betweenness centrality and VTE are not significantly different (p > 0.05). Despite they use different formalisms, these two indexes capture largely the same thing – namely, the amount of influence a node has over the flow of information in a graph, or over navigational paths. While it makes intuitive sense that a junction which is a strong conduit between locations (high betweenness) might require more deliberation, our analysis of RGI provides a further potential mechanistic explanation. We reveal that locations with a high information cost necessary to invest to reach the goal efficiently occur at such nodes and are associated with high levels of deliberation (35,45). VTE rates are correlated with another graph-theoretical index, closeness centrality (Pearson ρ = 0.44), which is defined for each node as the sum of the inverse of the length of the shortest path to all the other nodes. However, this correlation is significantly weaker than the correlation between RGI and VTE (p < 0.05). Unsurprisingly, VTE was higher at junctions than straight sections or corners (node degree centrality 2 vs. 3, Supp. Figure S3). We also report the novel finding that female participants were more likely than males to show VTE in task (Supp. Figure S4A). Task instruction had no impact on VTE (Supp. Figure S4B).

Finally, we compared the VTE values for participant who took the shortest path and those who took longer paths and found that with both the Go To Goal (KS=0.18, p<0.001) and the Take Shortcut (KS=0.19, p<0.001) instructions, the participants who chose the shortest path had a statistically significant lower value of VTE. This result suggests that participants who chose the shortest path could have planned it before starting moving; hence, they did not need to deliberate and engage in VTE behavior at decision points.

### Participants instructed to Take Shortcut select shorter routes but use more Control Information and are less optimal

The original study of Boone et al. (33) reported that participants who received the *“Take Shortcut”* instruction tended to select shortcuts more often than those who received the *“Go To Goal”* instruction and hence had a better performance, if this is measured only in terms of reward (solution length, see Figure 6A). Furthermore, the study of (33) investigated whether gender differences were present in the task and whether or not they are instruction-specific. The results of the study indicate a gender difference in strategy selection: both men and women taking more shortcuts when instructed to do so, but men were more likely to take shortcuts than females, under both kinds of instructions. However, it was not previously explored whether the increase of reward comes at the expense of an increase of Control Information and an overall decrease of optimality in the InfoRL sense (i.e., distance from the optimal Reward / Control Information curve).

**Figure 6:**
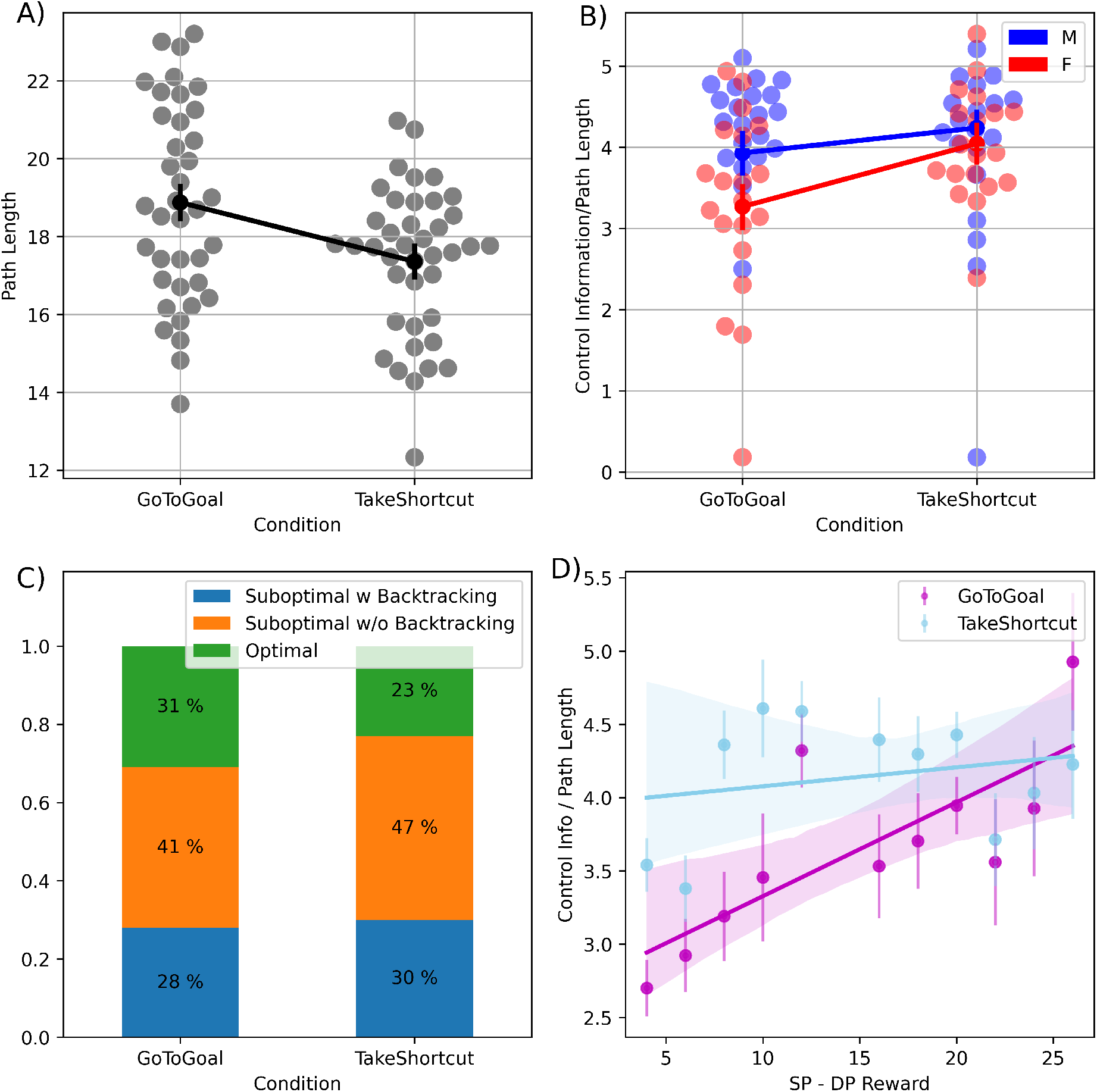
Group and gender differences in the amount of Control Information (divided by the number of steps of the solution) used in the task. (A) Difference in path length between “Go To Goal” or “Take Shortcut” instructions; black dots represent the means for the two conditions with respective standard errors as error bars, blue and orange swarm plot show the subject individual means. (B) Comparison of Control Information used by the two groups of participants who received “Go To Goal” or “Take Shortcut” instructions, split by gender (male or female); similarly to Figure 6A, green (male) and purple (female) dots represent the means for the two conditions with respective standard errors as error bars, blue and orange swarm plot show the subject individual means. (C) Distribution of the strategies used by the “Go To Goal” and “Take Shortcut” groups. Green: optimal strategy. Blue and orange: suboptimal strategies with backtracking and without backtracking, respectively. Please note that here the optimal strategies are those that lie on the optimal Reward / Control Information curve, not (necessarily) the shorter paths. (D) Amount Control Information invested as a function of the reward difference between the shortest route and the default policy. Each point represents the average Control Information invested by participants in the trials having a given reward difference. Error bars indicate standard errors. The two lines show the least-square fits for the two instructions: Go To Goal (magenta) and Take Shortcut (light blue). See the main text for explanation.

Here, we asked whether the instructions received by the two participant groups (*“Go To Goal”* vs. *“Take Shortcut”*) might have determined a different use of Control Information. We additionally looked at differences in Control Information between male and female participants, to assess whether this InfoRL measure captures the same gender differences reported in the original study (33). To investigate the main effect of Instruction (*“Go To Goal”* vs. *“Take Shortcut”*) on the amount of Control Information used by the participants and the interaction between Instruction and Gender (Male vs. Female), we performed a Mixed-Effect ANOVA with subjects as random effects. Note that in order to compare trials with different start-goal distance, in which participants (had to) take longer or shorter routes to the goal, here and later in the manuscript we divide Control Information by Path Length, or the length of the path selected by the participant from start to goal.

We found a main effect of Instruction, which implies that participants who received the instruction to Take Shortcut used more control information per step than participants who received the instruction to Go To Goal (TS - GTG = 0.52, t(77.48)=2.431, p=0.017). We also found a main effect of Gender, which implies that overall, male individuals show an increased Control Information per step compared to female participants (M - F = 0.44, t(86.46)=2.097, p=0.038). Furthermore, we considered an additive model in which both main effects are included and compared it with an interaction model that considers the interaction Instruction x Gender. The interaction model did not result in a significant reduction of the deviance with respect to the additive one (χ^2^(1)=0.8317, p=0.361). Results in accordance with the additive model showed that overall, male individuals use greater control information and that the increase in Control Information in the Take Shortcut condition is the same for both male and female gender groups (Figure 6B).

This pattern of our results provides a more subtle separation in the data not observed in (33). The authors reported that male participants tended to select shorter routes more often than female participants, which is in keeping with our finding that male participants use overall greater control information. However, the authors only reported a significant difference between the path length of the solutions selected by female (but not male) participants between Go To Goal and Take Shortcut instructions. Our more refined analysis, which is based on an information-theoretic quantity (Control Information) shows instead that both male and female participants are sensitive to task instructions and modulate their control information investment accordingly. Our analysis therefore highlights a novel result not evident from purely looking at path length.

Finally, we subdivided the strategies of the participants in the *“Go To Goal”* and *“Take Shortcut”* groups into Optimal strategies that lie on the optimal Reward / Control Information curve (green) and suboptimal strategies that lie under the optimal curve; see Figure 6C. We further subdivided the suboptimal strategies into those that include some backtracking (i.e., the participant passes twice on at least one node, *“Suboptimal with Backtracking”*, blue) or does not include backtracking (*“Suboptimal without Backtracking”*, orange). We found that participants who received the instruction to “Take Shortcut” selected less often the solutions that are optimal in terms of InfoRL compared to the participants who received the instruction to “Go To Goal” (log odds ratio = -0.3638, z=-4.259, p < 0.001). Notably this does not imply that the participants who received the instruction to “Take Shortcut” selected longer paths; indeed, as reported already in (33), the opposite is the case (Figure 6A). Rather, it implies that those who received the instruction to “Take Shortcut” were less optimal in the use of the (greater) resources that they invested. This result therefore highlights a difference between the InfoRL measure of optimality used here, which jointly considers path length and Control Information and the more limited measure of optimality used in (33), which only considers path length.

### Participants who receive the Go To Goal instruction modulate their resource investment depending on the benefits of finding the shortcut

To further investigate if participants modulated their usage of Control Information depending on task demands, we plotted Control Information as a function of the reward to be potentially earned by selecting the shortest route in each trial, i.e., the difference in reward between the shortest route and the default policy (Figure 6D). We made two separate linear least-squares regression analyses for the two instructions, since we hypothesized that instructions made a difference: indeed, the Go To Goal instruction leaves much more freedom in the choice of the Control Information compared to the Take Shortcut instruction. We reasoned that the participants who received the Go To Goal instruction were free to choose how much Control Information to invest; and they might have considered what was the reward difference between the shortest route and the default policy. In the trials where the reward difference was small and there was not too much to earn, they could have privileged a lower investment of Control Information, and vice versa for trials where the reward difference was greater. The results reported in Figure 6D confirm that participants who received the Go To Goal instruction modulate their Control Information depending on reward earnings; this is evident by considering that the slope of the magenta regression line is significantly different from zero (magenta line, slope = 0.06, p = 0.01). However, this does not happen for participants who received the Take Shortcut instruction (light blue line, slope = 0.01, p = 0.49).

### Participants vary their levels of Control Information and their (sub)optimality across trials

Finally, we tested whether the amount of Control Information deployed by each participant and the level of suboptimality (distance from the optimal Control Information per step / Reward curve) remain stable or change over trials. We reasoned that people might remain consistent across trials, showing a similar amount of Control Information and of (sub)optimality, i.e., distance from the optimal curve. However, our results show that people tend to vary their Control Information and (sub)optimality levels across trials, see (46) for a related result. Figure 7 shows that for most participants, the median value and variance of Control Information are highly variable across trials, with one exception being the (very few) participants shown at the beginning of the top panel, who always use very little Control Information, meaning that they invariably select the default policy when they receive a Go To Goal instruction. The same pattern of results emerges when considering the variance of the Distance from the optimal curve (Supp. Figure S5).

**Figure 7:**
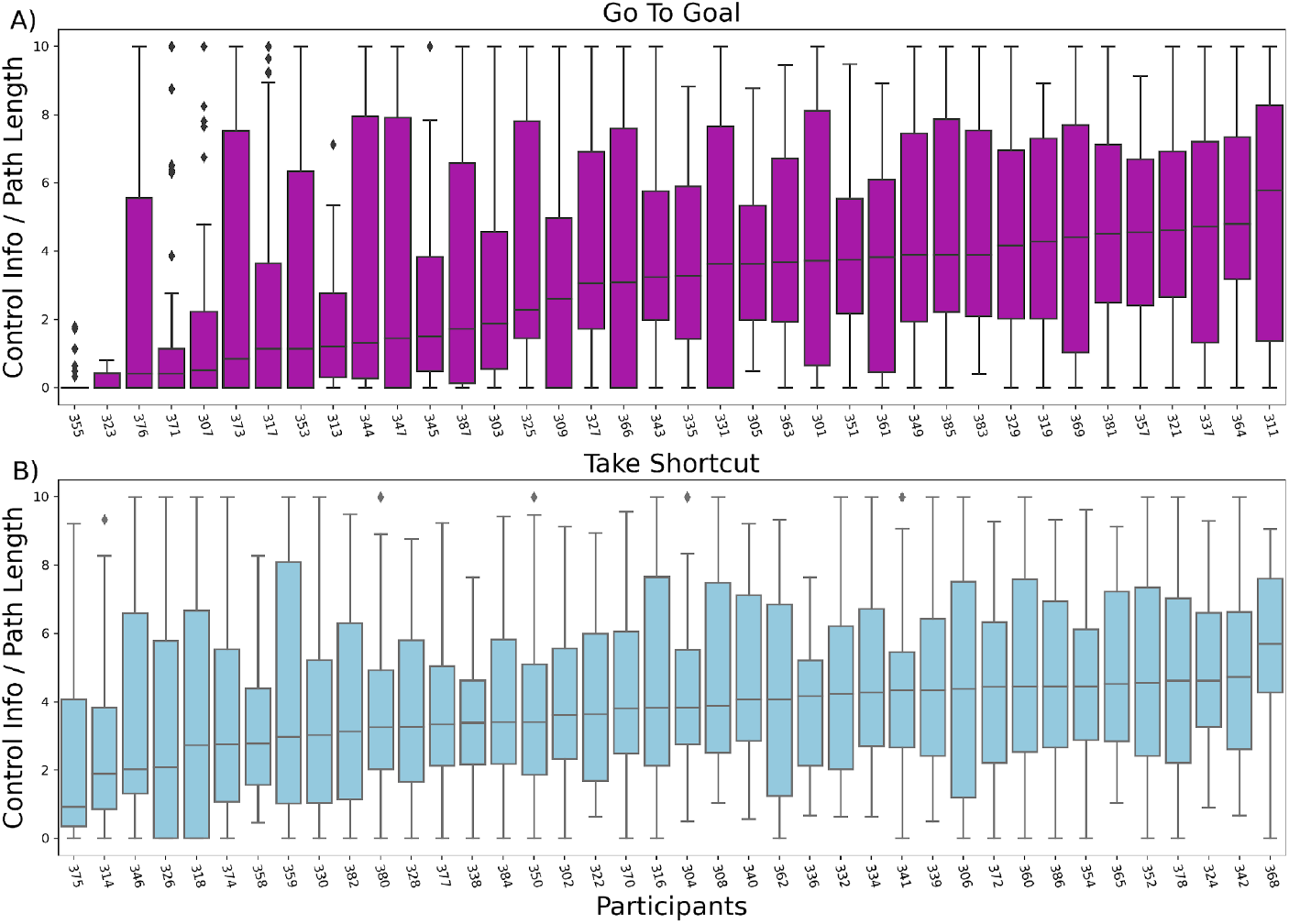
Boxplots of the Control Information divided by trial Path Length for each participant ordered by median value and subdivided by group (magenta: “Go To Goal” group, light blue: “Take Shortcut” group).

Given our finding that participants in the “Go To Goal” condition adapted their resource investment to the benefits of finding the shortcut (Figure 6D), we expected the variance to be instruction-specific. To assess this, we compared the variances of Control Information and Distance from the optimal curve, between the two groups of participants who received the instructions to “Go To Goal” and “Take Shortcut”, respectively. We found a main effect of Instruction in comparing the variance of Control Information (Levene’s test (*W* = 53.85, *p* = 2.82 * 10^−13^)), with the “Go To Goal” group having a larger variance in the two participant groups (see also Supp. Figure S6).

## Discussion

Here, we examined whether it is possible to formally characterize the choice of navigational plans in terms of a *bounded rational* process that trades off the quality of the plan (e.g., its length) and the cognitive cost required to implement it. We used the InfoRL scheme (29) to characterize formally the cognitive cost of devising a navigational plan, as the informational cost to deviate from a well-learned route (“default policy”). This formalism permits characterizing explicitly the trade-off between the costs and the benefits of devising a better plan, in terms of a Reward/Control Information curve that shows how much reward a participant can obtain by investing any level of control information. InfoRL therefore supports a richer notion of (bounded) optimality compared to the usual notion that the optimal behavior is the one that selects the shortest path. In InfoRL, “optimal” here refers to the possibility to obtain the maximum reward from the control information that one has invested, not the maximum level of available reward. In other words, InfoRL assumes that people are optimal if they get the maximum reward from the level of resource they invested (i.e., if they lie exactly on the optimal Reward / Control Information curve exemplified in Figure 1) and suboptimal otherwise. Using this definition, participants can be defined as optimal regardless of the information resources that they use, providing that reward they obtain is commensurate with their resource investment. For example, two participants who decided to invest few or many resources can be equally optimal (or suboptimal) if they lie on the optimal Reward / Control Information curve (or below it).

Our results show that InfoRL permits characterizing formally the different strategies that participants use during a navigational task (e.g., follow the well-learned route or take a shortcut). InfoRL also characterizes their associated information costs (e.g., very low for plans closer to the well-learned route, higher for shortcuts), and the optimality (or suboptimality) of the solutions, which correspond to the fact that participants achieve (or not achieve) the theoretically maximum reward for the amount of control information that they used. As shown in Figure 4, the theoretical Reward/Control Information curve of InfoRL retrieves nicely the set of navigational paths that people use during navigation in virtual reality (33). While optimal paths (at different levels of Control Information) lie exactly on the curve, other, suboptimal variants of the same paths can be readily identified, by considering that they lie diagonally to the corresponding optimal solutions. Furthermore, the results of (33) were obtained by manually labeling the routes taken by the participants, whereas here we used InfoRL to identify paths directly from data, without manual labeling.

Our analysis also shows that during navigation, participants exhibit more deliberative, “vicarious trial and error” (VTE) behavior (39,41) in the places of the maze where action selection requires using more information. While the notion of Control Information of InfoRL summarizes the information demand of planning an entire path, we used the (analogous) notion of Relevant Goal Information (37) to formalize the information requirement at each decision point. The strong correlation between Relevant Goal Information and a behavioral index of deliberation - VTE behavior - indicates two main things. First, they do not complete the whole plan in advance but rather continue deliberating and planning during the task; this is in keeping with a large body of evidence indicating that deliberation and action are intertwined during naturalistic behavior (42,44,47–51). Second, people invest cognitive resources preferentially in places in which information demands for action selection are greater, suggesting that they are sensitive to information demands of the current task (45,52,53). This latter result is in keeping with studies reporting that rodents exhibit greater VTE behavior at difficult decision points, in which information demands are greater - for example, when a conflict arises between different navigation strategies (54,55). In turn, VTE behavior might be instrumental to the evaluation of future choice outcomes, given that it has been associated with neurophysiological markers of deliberative information processing, such as hippocampal forward sweeps and transient representations of reward in the ventral striatum (56–58). It remains to be investigated in future studies to what extent the notion of Relevant Goal Information used here could be used to predict where VTE and hippocampal forward sweeps might occur more often on mazes.

Furthermore, our analysis shows that participants adapt their information investment to task demands, with greater investment for the Take Shortcut than the Go To Goal instruction, for both male and female participants. In this setup, the Take Shortcut instruction can be considered more challenging as it forces participants to significantly deviate from their default policy (which corresponds to a cost in InfoRL) and because shortcuts increase the likelihood of crossing the middle of the space, where the information required for action selection is greater (see Figure 5). The original study of (33) found a difference in the choice of strategies between conditions but only for female participants, whereas our refined analysis shows that also male participants modulated their level of Control Information in an instruction-sensitive way. It is also worth noting that participants who received the Go To Goal instruction could potentially use the default policy on every trial. However, they often select a shorter path, even if it costs more in terms of cognitive resources. This might be the case because they could consider minimizing other costs, such as the time spent to solve the task or the physical energy cost associated with getting to the goal, in addition to (or instead of) the cognitive cost. One indication that this might be the case comes from the finding that participants who received the Go To Goal instruction invested more Control Information when there was a greater difference between the rewards to be earned by following the shortest route versus the default policy. Future studies that manipulate multiple costs simultaneously might help shed light on the structure of preference of participants during navigation and wayfinding.

A separate analysis of the optimality of the paths shows that the more challenging “Take Shortcut” instruction leads to more suboptimality in strategy selection, see Figure 7B. In other words, while participants in the “Take Shortcut” group appear prima facie to be more effective (as they select on average shorter routes), our analysis shows that this comes at the cost of a greater investment of Control Information and - less obviously - this implies a greater risk of selecting suboptimal routes (i.e., routes that lie farther from the optimal Reward / Control Information curve). A possible explanation for this latter result is that there are many more ways to be suboptimal (in InfoRL terms) if one searches, remembers or follows the shortest path, than if one reuses a default policy. The difficulty of searching for an efficient solution is likely a key factor in making the instruction to Take Shortcut more challenging compared to Go To Goal instructions (25,59).

Finally, an additional analysis shows that people do not always use the same level of Control Information and do not always achieve the same (sub)optimality level, but tend to vary these two levels across trials, as shown also in other analyses of the same task (46). It is possible that this variability arises from moment-to-moment fluctuations in participants’ motivation to engage cognitive resources, from the fact that participants remember some parts of the map better than others, or (at least in part) as a byproduct of the adaptation of strategy selection to task rewards that we found in participants who received the “Go To Goal” instruction (Figure 6D). Testing these and other hypotheses for choice variability remains an open objective for future studies.

Taken together, the results of our InfoRL analysis nicely complement previous studies of the same virtual maze task that assessed reward (path length) maximization and the choice of navigation strategies as a function of instructions (33) and that used strategy-based path planning algorithms to simulate human trajectories and assess their variability (46). While each specific study used a different approach to model navigational strategies, there is a nice convergence of results; for example, the different analyses suggest that participants who receive the “Take Shortcut” instruction tend to choose shorter routes, that they select not just optimal routes but also suboptimal routes (33) and that they show a high variability in their strategy selection (46). This convergence is reassuring and supports the usefulness of InfoRL to model spatial navigation tasks, as a complement to the methods exemplified in the above studies. Moreover, some of our results highlight the usefulness of a computational approach such as InfoRL that considers both the costs and the benefits of strategy selection, rather than a more naive assessment of navigation efficacy, such as path length. Describing navigational planning and wayfinding as bounded rational processes permits going beyond a narrow notion of resource-independent optimality as “maximizing reward” and consider to what extent participants are able to optimize their performance, relative to the amount of information that they process (52,60–63). For example, our analyses show that about a third of the participants lie very closely to the optimal curve described by InfoRL in the less challenging (Go To Goal) condition, but only around a quarter did so in the more challenging (Take Shortcut) condition. This result emerges despite participants who receive the Take Shortcut instruction select (as expected) shorter routes and would be therefore considered more optimal in reward-maximization terms. Furthermore, our finding that participants deliberate more in places where information demands are greater are in keeping with previous studies of human vicarious trial and error (40,64) and suggest that people are sensitive to information costs and allocate them adaptively - which in turn lends support to information-theoretic analyses of planning and deliberation (26,27,53). Our finding that participants who receive the “Go To Goal” instruction (and are therefore more free in their strategy selection) change their resource investment depending on the benefits of finding the shortcut provides converging evidence for the importance of adaptive allocation of limited resources and offers a potential explanation for some of the variability that is found in participants’ behavior. Our finding that female participants engage in more deliberation is novel and may relate to greater spatial anxiety on average in females compared to males (65). Further research exploring this will be useful.

InfoRL and related approaches based on mutual information, free energy or similar measures have been widely investigated in theoretical studies and to address simple decision or control settings (19,23–26,45,61,66–75), but not to study more complex behaviors such as wayfinding as done here. One of the reasons is that formalizing (default) policies for planning can be challenging. In theoretical studies, default policies are assumed to be random, but random behavior is not plausible during navigation. Here, we defined the default policy as a well-learned route, but this might not be possible in other studies that do not include an initial training phase. Previous studies suggest that during wayfinding, people might consider a default “vector-based” strategy (1) or follow the available affordances (76,77). Specifically, line of sight along different options at decision points may be a strong affordance based feature driving a default policy (78). Vector-based, affordance-based and other strategies could be potentially incorporated as “default policies” in InfoRL but the efficacy of these methods to characterize human behavior and cost-benefit computations remains to be tested.

The results reported in the current study depend on at least two assumptions: 1) that the participants’ default policy is the habitual path extensively practiced before the experiment, 2) path length is a suitable measure of value and reward. With regard to the default policy, while this seems reasonable in this setup, one side effect is that following the default policy backward entails a high amount of Control Information, which is not necessarily plausible (and could be relaxed in future analyses). More broadly, the choice of a default policy is a critical decision for any InfoRL study. Most simulative studies use the random policy as the default policy (22,24), this might be less compelling in the case of human or animal navigation studies. Intriguingly, a rodent study suggests that the default behavior of animals is much closer to an optimal policy for foraging than a random policy (30) and also human navigation follows regularities (1) that might be considered as a “default” in an InfoRL setting. Furthermore, one can find regularities not just in human or animal behavior but also in the way cities are designed, e.g., in such a way that they afford an easy navigation to salient locations. This is somewhat reflected also in the design of the experiment of (36), in which the habitual route is at the periphery whereas shortcuts are almost always in the middle. This is a sensible design choice, since if the default were through the center of the maze, then it would be very difficult to obtain a large number of start-goal configurations in which a shortcut (passing through the periphery) exists. In turn, a paucity of shortcuts would “flatten” the theoretical InfoRL curves shown in Figures 1B and 4A, since selecting a path that differs from the default policy would not increase reward. At the same time, the choice to place the habitual route at the periphery and the shortcuts at the middle implies that shortcuts are invariably linked to places having high relevant goal information or betweenness centrality. Future experiments could therefore consider different designs in which shortcuts and places with high relevant goal information or betweenness centrality could be disentangled more clearly.

With regard to the use of path length as a measure of value and reward, it is possible that other metrics could be explored, such as time taken, smoothness of paths, angular displacement from the goal. Such other metrics have been successfully explored in studies examining behavior in humans and rats in relation to RL (79,80). Such metrics could be incorporated into future studies, particularly where these vary, such as open mazes and mazes with dead-ends (78). In addition to analysis of VTE it would also be useful to explore eye-movements and neural activity in future studies with an InfoRL approach. Eye-movement analysis combined with RL-agent modeling have proved useful in exploring the underlying mechanism involved in navigation (81). For human imaging or neural recording studies it may be possible to probe the responses with regressors such as the amount of Control Information alongside various other regressors explored in studies of navigation (11,82).

This study has some limitations that could be addressed in future studies. First, it is worth noting that InfoRL speaks to a specific notion of effort and decision complexity that does not fully cover other sources of difficulty that arise during decision-making. The instruction to “Take Shortcut” is *more challenging* in InfoRL terms, since it requires investing cognitive effort (27); however, it is also *less ambiguous*, as it specifies that a shortcut is the only valid solution. On the contrary, the instruction to “Go To Goal” is more ambiguous as it allows for several solutions, at different levels of cognitive effort (e.g., do I simply re-activate the learned path or try a risky new one?) The InfoRL framework does not fully capture this notion of ambiguity, which would require more sophisticated methods. Another limitation of the study is that the design of the experiment does not fully separate information- and graph-theoretic measures of relevant goal information, betweenness centrality and closeness centrality, all of which are correlated. Future studies could consider maze designs that better disambiguate between these measures and permit assessing their effects on navigational planning (83)). Furthermore, it would be possible to design mazes in which the costs of shortcuts vary, or adapt navigation to hunting (84), to study how these costs influence the participants’ propensity to select shortcuts or default policies.

## Methods

### Markov Decision Processes

A Markov Decision Process (MDP) is a mathematical framework for modeling decision making in aleatory environments. It is defined by a tuple ⟨*S, A, R, P, π*⟩ where: *S* {*s*_1_, …, *s_N_*} and *A* = {*a*_1_, …, *a_M_*} are, respectively, finite sets of N states and M actions; *R* = *R* (*a, s*) is a scalar function representing the reward obtained in state *s* after taking action a; *P* = *P* (*s*′|*s, a*) is a transition model obeying the Markov property - i.e., the outcome for taking action *a* in state *s* depends solely on the current state *s*, and not on the history of states and actions that precedes it -, where *P* (*s*′|*s,a*) represents the probability of transition to state *s*′ when taking action *a* in state *s*; and *π* = *π*(*a*|*s*) is a state to action mapping, which represents the stationary probability for the agent to take action *a* in state *s*, and is known as the agent’s *policy*.

In this article, we focus on episodic pathfinding tasks, i.e., tasks where the aim is to reach a terminal state *s*_0_, with the absorbing property *P*(*s*_*goal*_|*s*_*goal*,_ *a*) = 1 ∀*a* ∈ *A*. Additionally, the reward function is negative for all states and actions combinations outside the absorbing state and zero otherwise: *R*(*s, a*) < 0 ∀*a* ∈ *A, s* ∈ *S*, and *R*(*s*_*goal*_, *a*) = 0 ∀ *a*.

We define the value function of a policy *π* as the expected accumulated reward for executing *π*starting from state *s*_0_:

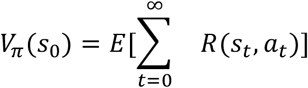

where the expectation is taken over the probability of all future trajectories, starting in *s*_0_ and executing *π*, with *Vπ*(*s*_*goal*_) = 0.

Furthermore, we define the optimal value function, 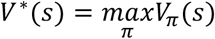, as the maximal *π* achievable value, and we define optimal policy *π*\* the policy that achieves it.

### Control Information

A starting point of InfoRL is that searching for the shortest path (or optimal policy) requires a cognitive or computational effort. Thus, next to the concept of value, we introduce the concept of Control Information (or control cost) to deviate from a default or habitual policy.

We define the Control Information to execute a policy *π* in state *s*, with regard to a policy *ρ*, as the Kullback-Leibler divergence between the two policies in state s:

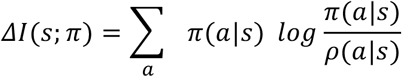

where *ΔI*(*sπ*) = 0 for terminal states, and *ρ* is an arbitrary “default policy” which represents a standard behavior the agent follows when not exerting control. The Control Information quantifies in natural units (or bits, if the logarithm has base 2) how much the agent’s policy * deviates from the default policy *ρ* at a given state *s*.

Although the default policy is arbitrary, a common choice is to take it uniformly distributed over all the available actions in state *s* (24,22,23). In this work, instead, we define the default policy as the policy the participants learned during the training phase of the experiment.

In the previous section, we defined a value function for a policy as the expected accumulated reward for executing *π* starting from state *s*_0_. Here, following (22), we define a Control Information function as the expected Control Information cost over all the trajectories from a starting state *s*_0_, using a policy *π* and default policy *ρ*:

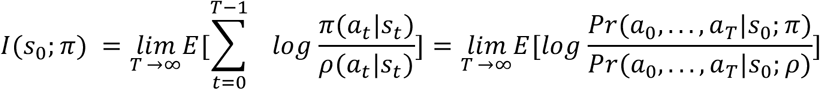

where the expectation is taken with respect to all possible paths starting in *s*_0_.

However, in our experiment, this formulation poses three issues. First, it requires assuming a default policy; as explained above, we assume that the default policy is the policy that follows the training path.

Second, unlike in simulation studies, we do not have direct access to the participants’ true policies policy *π*(*a*|*s*) for a given start-goal setup. However we can infer an approximation 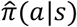 from the state-action sequence {*s*_0,_*a*_0,_ … *a*_*T* −1,_, *s*_*T*_} that corresponds to the paths selected by the participants, by computing a matrix of co-occurrences of states and actions 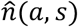. By normalizing this matrix, we obtain the joint probability of actions and states 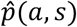. Consequently, we write the participant’s observed policy as 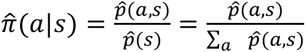, where 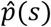 is the marginal probability of the states. Note that the joint probability is computed over a single participant’s trial trajectory, and as such the observed policy reflects a particular start-goal setup.

Third, unlike in simulation studies, in this experiment we cannot average across multiple trajectories to calculate Control Information, because participants completed single start-goal trials. Thus, we assumed this single sample to be representative of the statistics of the path distribution for the agent (as a unimodal distribution sharply peaked around the mean). This allows to define an “empirical” control information as:

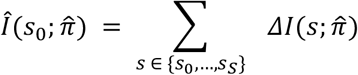

In the special case of a deterministic policy π(*a*|*s*), this is not a problem, because there is only a single path starting from *s*_0_ with nonzero probability, and (for the states visited by the agent) the inferred policy 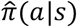 is exactly the agent’s policy *π*(*a*|*s*).

### Free energy formulation and InfoRL

By combining the definitions of value and cost from previous sections, in this section we expand the notion of optimality by taking in consideration not only the expected reward, but the expected Control Information cost exerted by the agent. This leads to changes in the shape of the value-information curve.

Following Rubin et al. (22), we define a free energy function *F*(*s*_0_ ; *β*) which combines the value and the information terms:

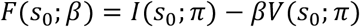

where the parameter *β* ≥ 0 controls the tradeoff between information and value.

We want to find the policy that maximizes the value, while keeping the information cost under a threshold:

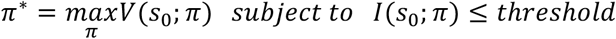

where the threshold on information cost can be thought of as a cognitive resource limit. This constrained optimization problem is similar to the one in Rate-Distortion Theory (85) and the Information Bottleneck (86).

Rubin et al. (22) showed that the optimal free energy vector *F*\*(*s*_0_; *β*) satisfies Bellman’s equation (87) and then they proposed an algorithm to calculate the optimal free energy vector, InfoRL, for a given MDP model and tradeoff parameter *β*.

Note that the solutions for different values of *β* are shown to form in the reward-information control domain a nonlinear concave curve (see Figure 1), whose shape is dependent on the topology of the map, the MDP model and which default policy is used. The points along the curve include all the optimal solutions under the free energy definition. The leftmost point (*β* → 0) represents the highest rewarding among the zero control information solutions (i.e., the agents followed blindly the default policy) while the rightmost (*β* → ∞) has the lowest control cost among the solutions with the highest reward (i.e., a shortest path strategy). In general, the solutions along the curve are the only solutions in the map showing an optimal resource (information control and reward) allocation strategy. In fact, the top-left area of the graph is unachievable (e.g.: solutions with higher reward than the shortest path). The bottom-right area includes all the suboptimal solutions, i.e., the agent pays a higher control information cost to obtain an equivalent reward to the optimal curve (shifted to the right of the optimal solution), or, equivalently, the agent achieves a lower reward for the control information cost paid, compared to the optimal curve. In practice, this means that suboptimal agents pay an information control cost for deviating from the default policy but spend it in a worse way than their optimal counterparts.

### Relevant Goal Information

The Relevant Goal Information (RGI) (35) is defined as the conditional mutual information between current goal and action given the current state. It is a way of quantifying how much information (in bits or nats) is required in a given state to reach a goal through action. This amount of information depends on task constraints (i.e., what kind of optimality is required). For example, let’s consider a navigational task in a two-room maze, where the goal consists in reaching a specific goal location with the shortest path (optimality condition). Let’s imagine we start from the bottom of one room, and we are told that the goal is in the left corner of the other room. For the first steps, the only information we need to keep in mind is that the goal location is in the other room, but not the specific goal location; this is because, regardless of the goal position, the action we must select is reaching the other room. Thus, at the bottom of the start room, the relevant goal information is low. However, when we enter the other room, we need to choose more carefully which direction to take to reach the left corner. Because knowing precisely the goal location is key for action selection at the entrance of the goal room, the relevant goal information is high.

Formally, the RGI can be defined as the difference (in bits) of the entropy of the action distribution at each state between the case mentioned above (when the goal location is known) and the case in which the goal location is unknown, and hence one cannot discard any action a-priori. As in our analyses we compare RGI with graph-theoretic measures (betweenness and closeness centrality), we use RGI *per state*, defined as follows:

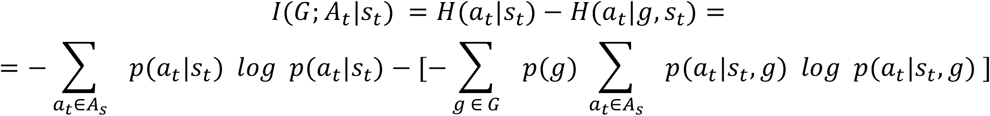

where *H*(· | ·) represents a conditional entropy between two variables. The RGI per state can be obtained using the Blahut-Arimoto (88,89) optimization algorithm.

### Vicarious Trial and Error

Vicarious Trial and Error (VTE) describes a stereotypical behavior that humans and rodents exhibit when uncertain at decision points (e.g., in T-mazes) and which consists in moving the head back and forth between the possible paths (39,41). This behavior has been shown to be associated with endogenously generated hippocampal dynamics called “forward sweeps” and the sequential activation of sequences of place cells that correspond to possible future locations (90).

Here, we measure a “*VTE event*” by considering when the participant halts (speed under a threshold *ν*_*threshold*_ = 1[*frame*^−1^]) and looks around (angular velocity over a threshold 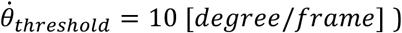, and “*VTE rate*” is the percentage of time over the entire trial time spent doing VTE.

## Acknowledgements

This research received funding from the European Union’s Horizon 2020 Framework Programme for Research and Innovation under the Specific Grant Agreement No. 945539 (Human Brain Project SGA3) to GP and the European Research Council under the Grant Agreement No. 820213 (ThinkAhead). We would like to thank Prof. Mary Hegarty and Dr. Alexander Boone for making their experimental data available.

## Data Availability Statement

The behavioral data used here are from Experiment 2 in Boone et al. 2019 and they have been made available by the study authors (https://osf.io/ykxts/). All the materials used for the analyses and the figures of this paper are freely available in: https://github.com/gllancia/Humans-account-for-cognitive-costs-when-finding-shortcuts

## Supplementary Materials

### Supplementary Video

The Supplementary Video V1 shows the different families of solutions discussed in Figure 4 of the main text.

### Supplementary Figures

**Figure S1:**
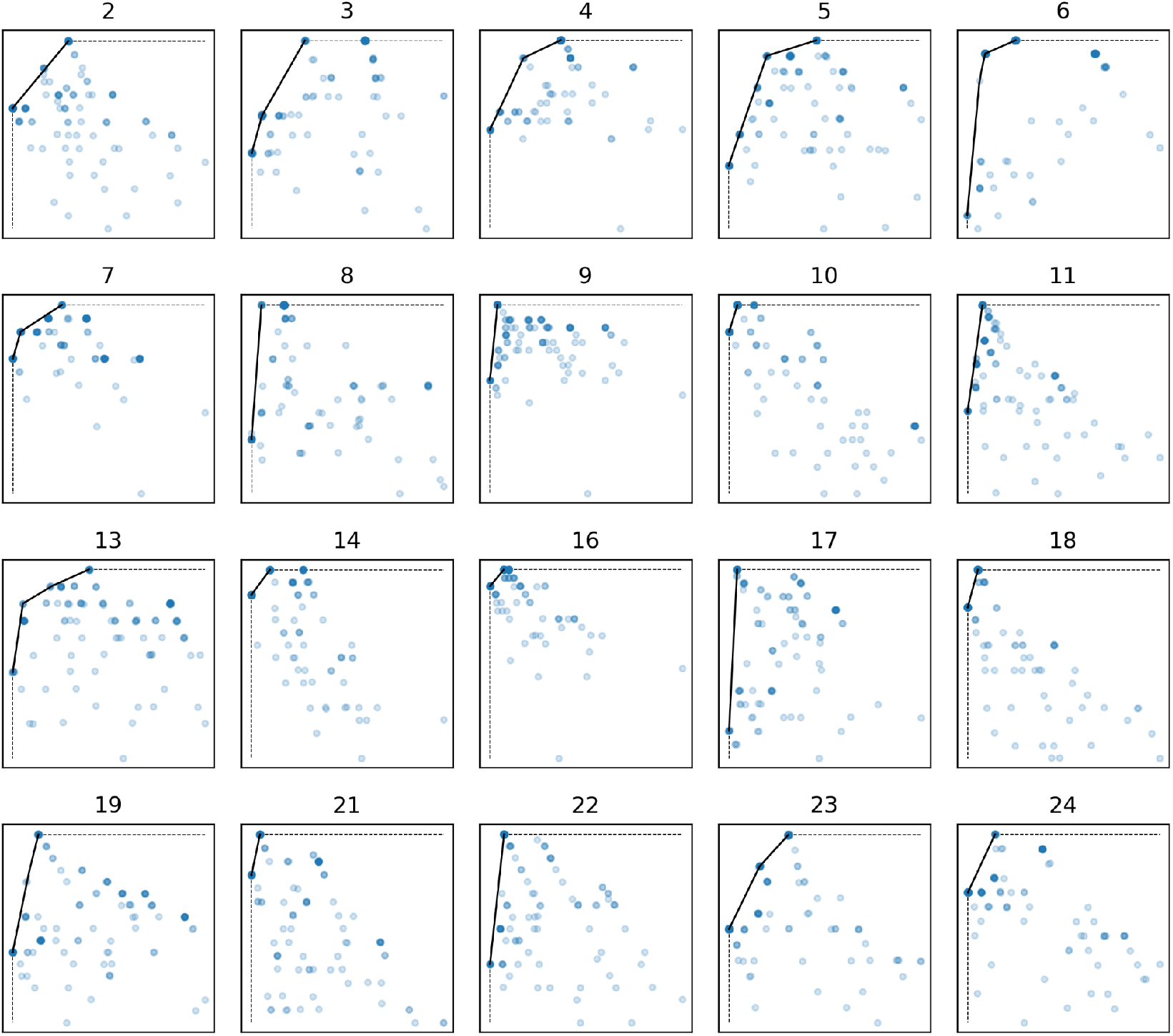
Reward / Control Information plots for the 20 pairs of start-goal locations collected by (33). The blue points represent participants’ single trial data (darker blue means more points overlapping), solid black curve is the optimal curve, and together with the dashed black curve delimits the unachievable area (upper left portion of the plot).

**Figure S2:**
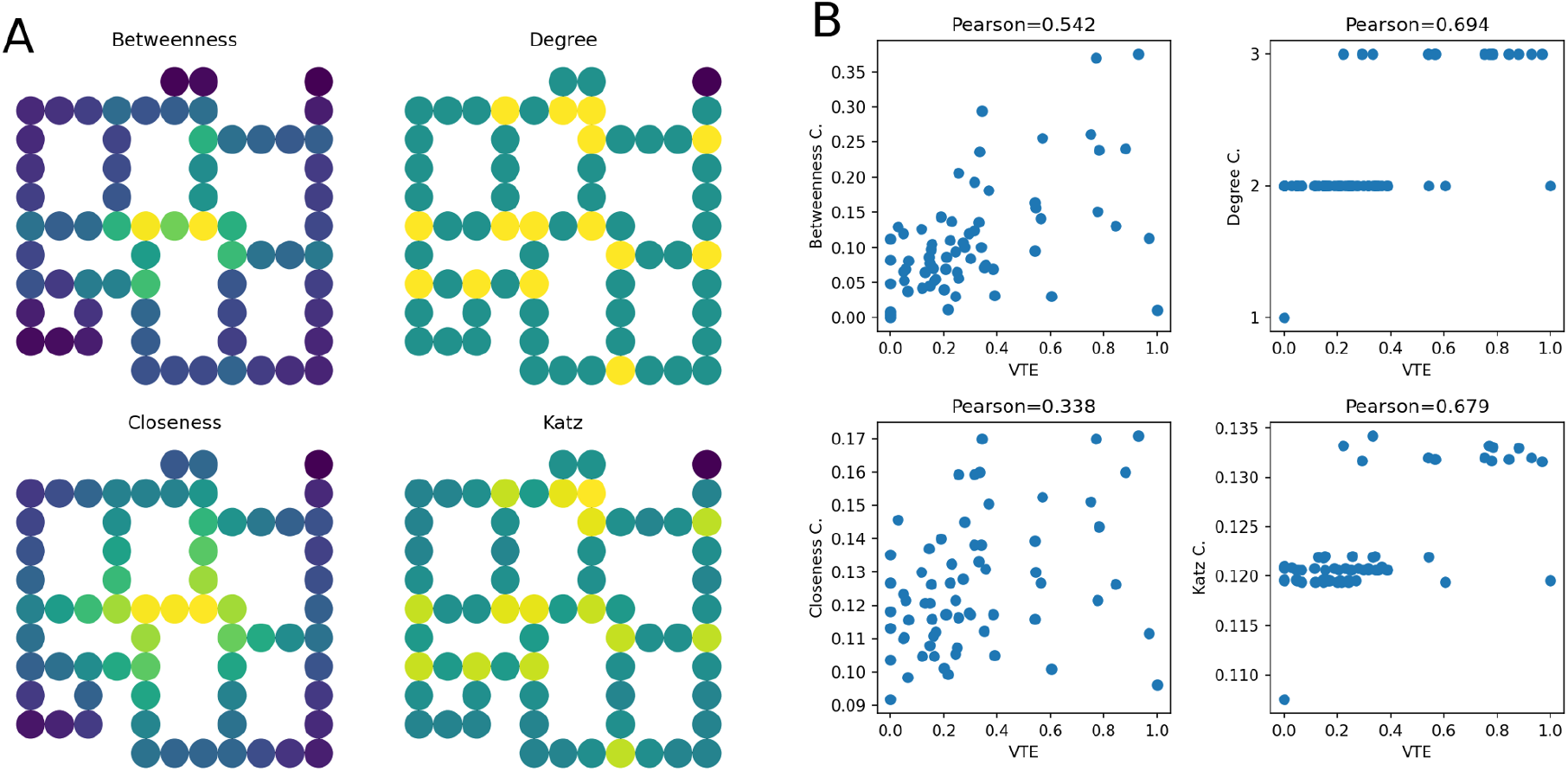
A) Betweenness, Degree, Closeness and Katz Centrality values for each state of the map; B) (Betweenness, Degree, Closeness and Katz) centrality value vs VTE value plots.

**Figure S3:**
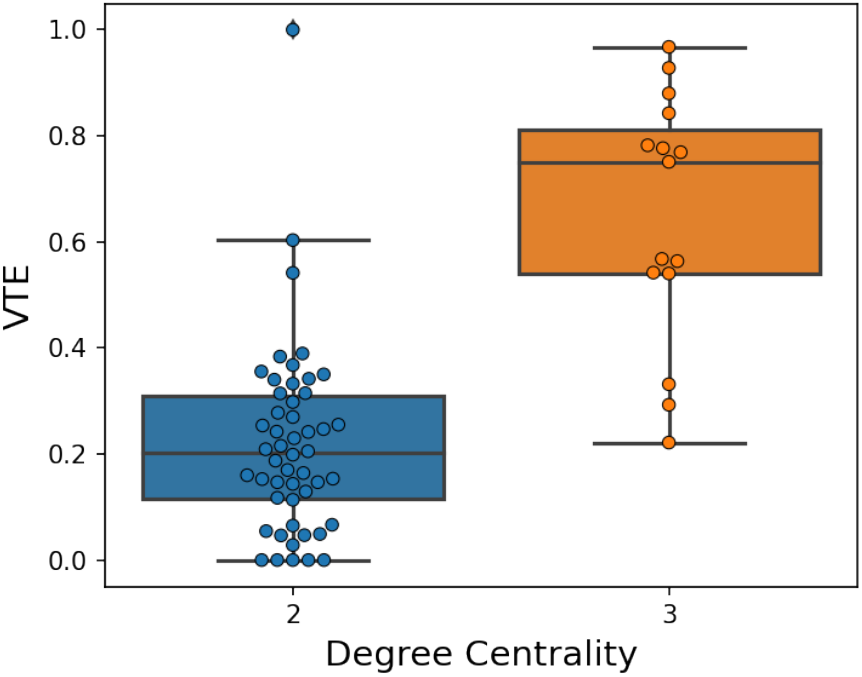
Scaled VTE for locations in the maze comparison for states with Degree Centrality equal to 2 and 3. A 2-sample Kolmogorov-Smirnov test confirmed (D=0.74, p<0.001) that states with Degree Centrality equal to 3 show greater VTE values.

**Figure S4:**
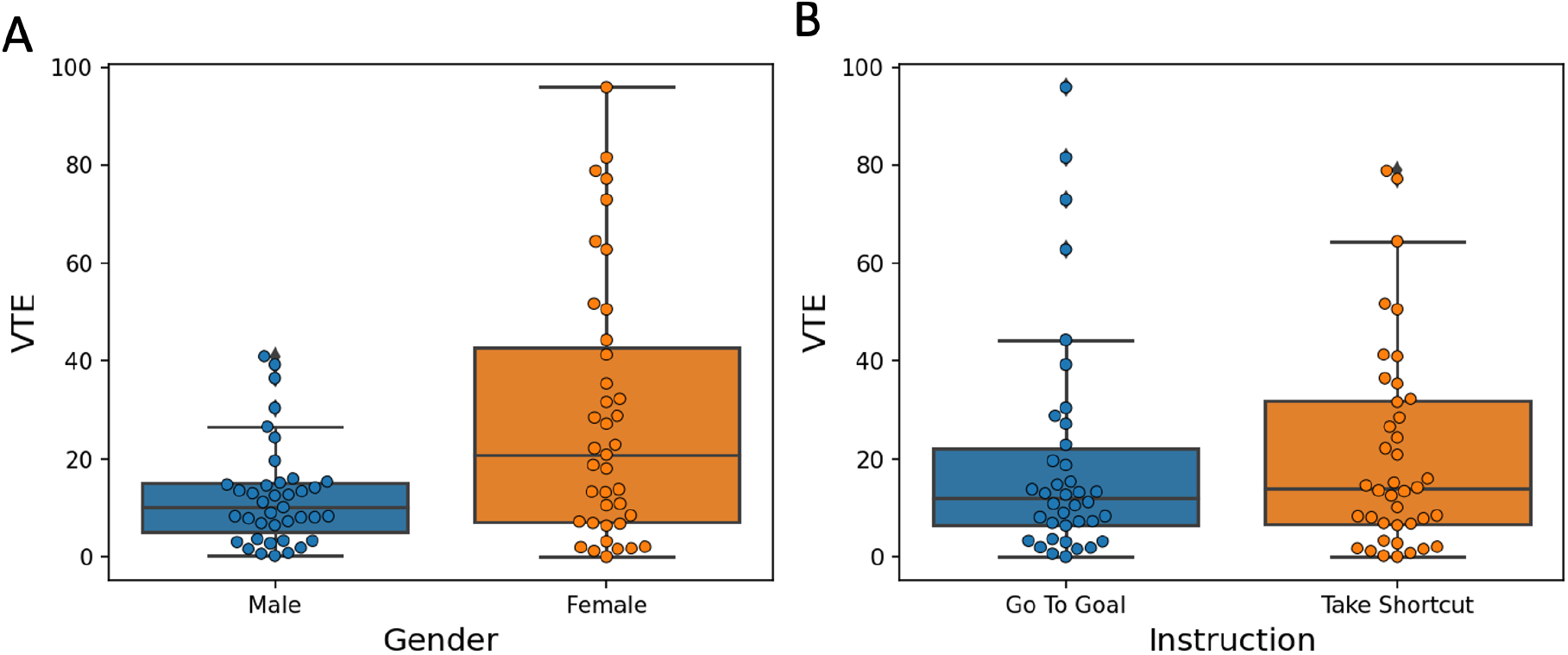
Left: Scaled VTE for locations in the maze compared by gender (D=0.38, p=0.006). Right: VTE values by condition (D=0.16, p=0.66).

**Figure S5:**
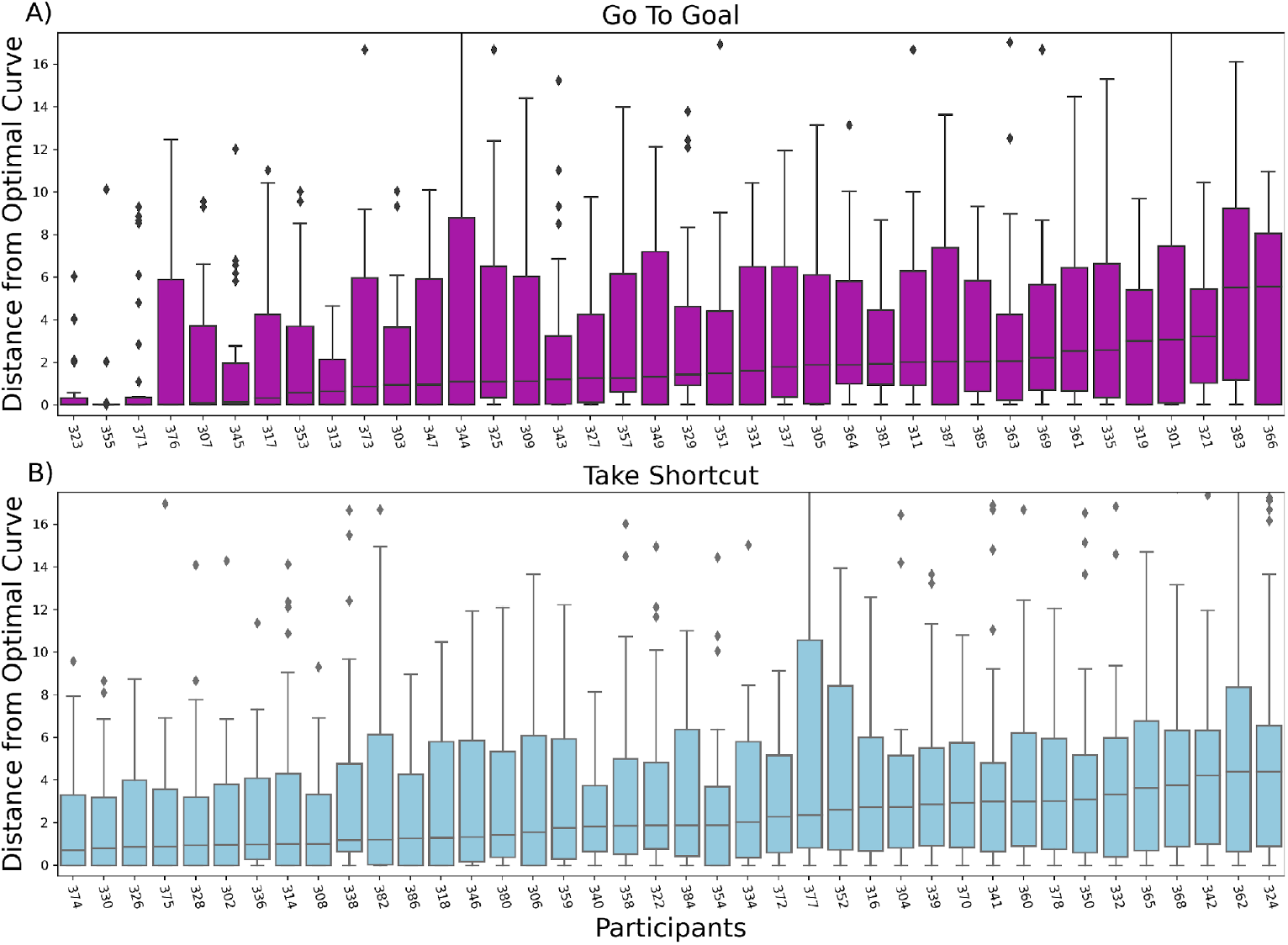
Boxplots of the Distance from the optimal curve (bottom two panels), ordered by median value and subdivided by group (magenta: “Go To Goal” group, light blue: “Take Shortcut” group); we found no significant difference in the variance of Distance from the optimal curve (W = 0.050, p = 0.82) in the two participant groups.

**Figure S6:**
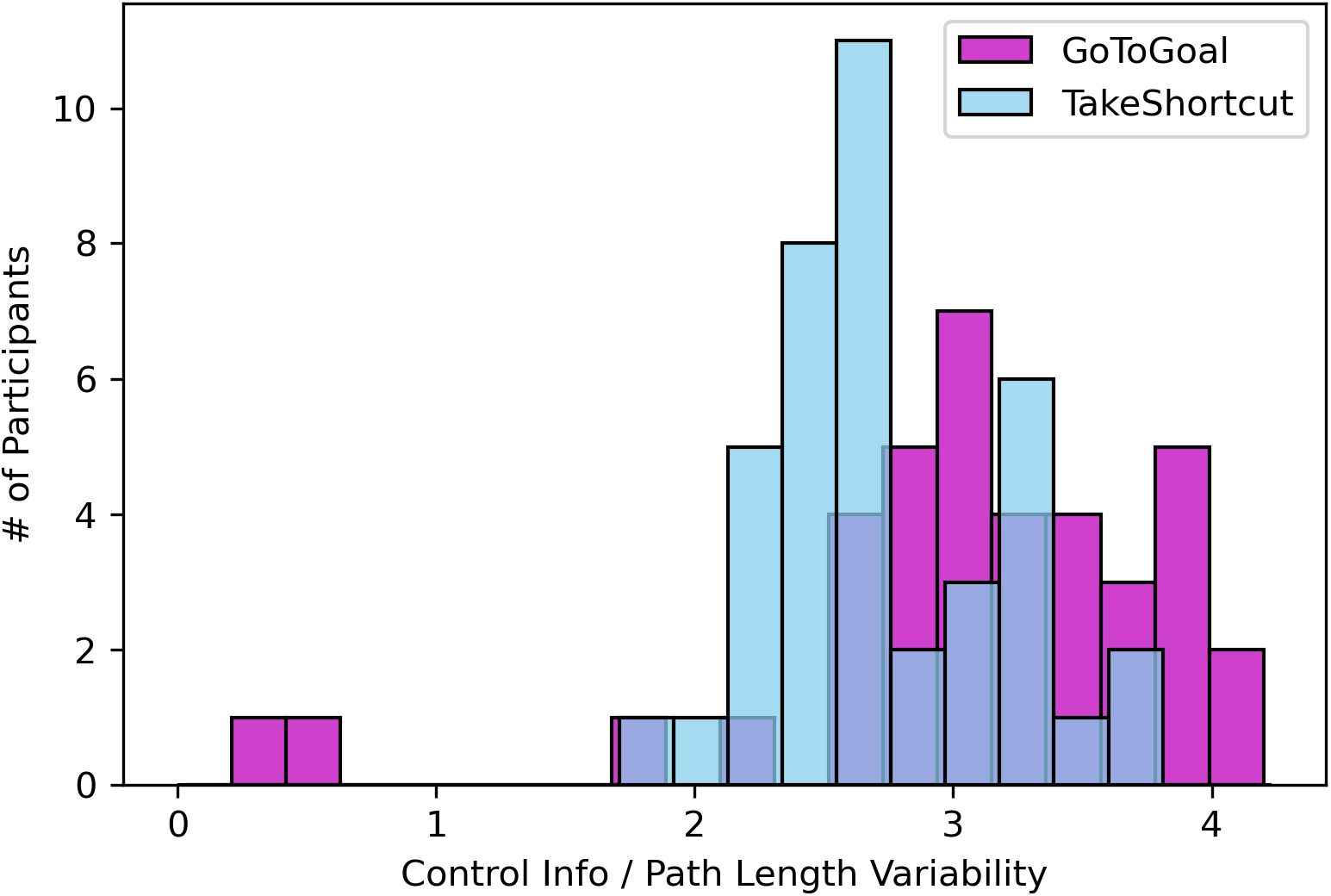
Histogram of the variability (i.e., standard deviation) of the Control Information – Path Length ratio across participants, as shown in Figure 7 (magenta: “Go To Goal” instruction, light blue: “Take Shortcut” instruction).

## Notes

### Competing Interest Statement

The authors have declared no competing interest.

### Summary of Updates

-

